# Consequences of the constitutive NOX2 activity in living cells: cytosol acidification, apoptosis, and localized lipid peroxidation

**DOI:** 10.1101/2021.02.23.429648

**Authors:** Hana Valenta, Sophie Dupré-Crochet, Tania Bizouarn, Laura Baciou, Oliver Nüsse, Ariane Deniset-Besseau, Marie Erard

## Abstract

The phagocyte NADPH oxidase (NOX2) is a key enzyme of the innate immune system generating superoxide anions (O_2_^•−^), precursors of reactive oxygen species. The NOX2 protein complex is composed of six subunits: two membrane proteins (gp91^phox^ and p22^phox^) forming the catalytic core, three cytosolic proteins (p67^phox^, p47^phox^ and p40^phox^) and a small GTPase Rac. The sophisticated activation mechanism of the NADPH oxidase relies on the assembly of cytosolic subunits with the membrane-bound components. A chimeric protein, called ‘Trimera’, composed of the essential domains of the cytosolic proteins p47^phox^ (aa 1-286), p67^phox^ (aa 1-212) and full-length Rac1Q61L, enables a constitutive and robust NOX2 activity in cells without the need of any stimulus. We employed Trimera as a single activating protein of the phagocyte NADPH oxidase in living cells and examined the consequences on the cell physiology of this continuous and long-term NOX activity. We showed that the sustained high level of NOX activity causes acidification of the intracellular pH, triggers apoptosis and leads to local peroxidation of lipids in the membrane. These local damages to the membrane correlate with the strong tendency of the Trimera to clusterize in the plasma membrane observed by FRET-FLIM microscopy.

**Highlights:**

- Trimera is a tool to trigger a continuous ROS production in living cells
- Continuous NOX2 activity causes cytosol acidification and apoptosis
- ROS overproduction leads to localized oxidation of the membrane lipids
- Trimera tends to clusterize in the plasma membrane of COSNOX and COS-7 cells

## INTRODUCTION

The NOX family of NADPH oxidases generates superoxide anion (O_2_^•−^) that is the precursor of other reactive oxygen species (ROS) [1]. Among them, the phagocyte NADPH oxidase (NOX2), is the best known and a prototype for other family members. It is a key player of the most effective mechanisms for the destruction of pathogenic microorganisms by the ROS production. This NADPH oxidase is a protein complex composed of six subunits; two membrane proteins (gp91^phox^ and p22^phox^) forming the catalytic core, three regulatory proteins (p67^phox^, p47^phox^ and p40^phox^) forming a ternary complex and a small GTPase Rac [1]. Lack of the NOX2 activity leads to chronic granulomatous disease (CGD) characterized by severe and recurrent infections. On the other hand, ROS overproduction contributes to cardiovascular and neurodegenerative diseases. Thus, the NOX2 activity needs to be finely regulated in order to produce the ROS where and when they are needed and in an appropriate concentration; either as antimicrobial effectors or as signaling molecules [2, 3]. The activity of the NADPH oxidase is controlled by the spatial separation of its membrane components from the cytosolic subunits. Upon activation either by phagocytosis or by a soluble stimulus such as phorbol ester (PMA) or arachidonic acid, the cell signaling cascade triggers phosphorylations of the ternary complex of cytosolic subunits. Those phosphorylations change dramatically its conformation into an active state able to interact with the membrane. Then the ternary complex migrates to the membrane independently of Rac. When it reaches the membrane, it associates with gp91^phox^, p22^phox^ and Rac-GTP forming the active complex able to produce superoxide anions. The formation of the active complex involves the reorganization of the intramolecular interactions within the cytosolic complex and formation of new interactions with the membrane subunits as well as with membrane lipids [4].

Berdichevsky *et al.* designed fusion proteins of selected regions of cytosolic proteins p67^phox^ and p47^phox^ with Rac [5]. The chimeric protein called Trimera is composed of the PX domain and the two SH3 domains of p47^phox^ (aa 1-286), the TPR domains and the activation domain of p67^phox^ (aa 1-212) and the full-length Rac1 containing the Q61L mutation assuring that Rac1 is in the active GTP-bound form (Figure 1) [5]. We recently observed that such a chimeric protein composed of the domains of cytosolic subunits, which are essential for the NADPH oxidase activity, turns the NADPH oxidase into a constitutively active enzyme in live cells [6]. The Trimera offers the opportunity to investigate the NADPH oxidase activity independently from any stimulation of the cellular signaling cascades. It is thus possible to focus on the active state of the NADPH oxidase and the consequences of long-term ROS production at the molecular level.

**Figure 1.**
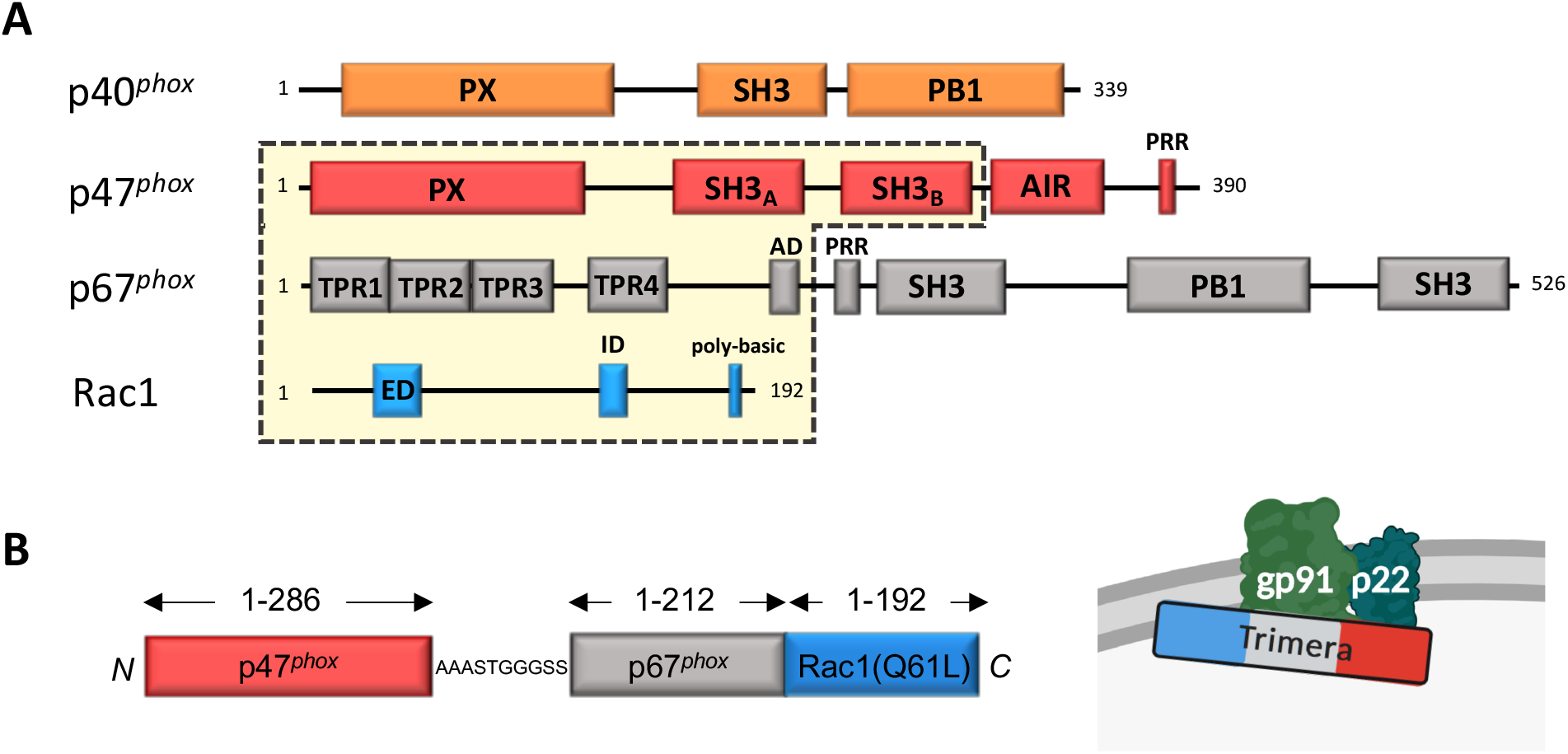
**A: Modular structure of the cytosolic subunits and Rac1.** The modules (domains) are represented by boxes whose size is proportional to the number of involved amino acid residues. AD is the activation domain, PRR is the prolin rich region, ED is the effector domain, ID is the insert domain. Domains in the yellow rectangle are conserved in the Trimera. **B: Chimeric protein “Trimera”.** Left: Trimera is a fusion protein created from the functionally important domains of p47^phox^, p67^phox^ and Rac1 (showed in A). Rac1 contains the point mutation Q61L, which keeps it in a GTP-bound form. AAASTGGGSS is a 10-amino acid linker between the p47^phox^ and p67^phox^ segments. There is no linker between p67^phox^ and Rac1. Right: Trimera can create the active NOX complex with gp91^phox^ and p22^phox^. Trimera is a single activating protein that can replace separated cytosolic proteins in the NOX activation process.

In this article, we describe the localization and distribution of the Trimera at the subcellular level in the cell membrane using fluorescent protein labeled Trimera. We likewise evaluated the consequences of the continuous and long-term NADPH oxidase activity on cell physiology in terms of cell viability, intracellular pH and lipid peroxidation.

## RESULTS

### Trimera enables the active state of the NADPH oxidase in cells

We expressed either the separated cytosolic subunits, p67^phox^, p47^phox^ and p40^phox^, each labeled with a fluorescent protein (FP) or the Citrine-Trimera [6, 7] in COSNOX cells, COS7 cells stably expressing p22^phox^ and gp91^phox^ subunits [8]. The COSNOX cells contain endogenous Rac1 [9]. The ROS production was detected by the L-012/Horseradish Peroxidase (HRP) luminometry assay (Figure 2). The COSNOX cells expressing the separated subunits were stimulated by phorbol ester, PMA. The PMA is a soluble stimulus, which launches phosphorylation of NOX subunits through activation of protein kinase C. As expected with this activation process, the luminescence signal raised slowly and reached a plateau at around 30 min. We also observed that p40^phox^ was not necessary to induce a strong NADPH oxidase activity and the presence of p40^phox^ actually decreased the ROS production almost twice (Figure 2A). This observation supports the hypothesis of p40^phox^ being a negative regulator in a PMA-stimulated situation [10, 11]. In COSNOX cells expressing the Trimera, a strong signal is detected immediately after the addition of the L-012/HRP mixture without any chemical stimulation. The signal raised quickly for a few minutes, likely the time necessary to reach the steady state of the detection reaction, and then reached a plateau (Figure 2B). This plateau can be seen as a “snapshot” in the time period during which the ROS are produced continually in presence of the Trimera. These results confirmed the previous observations of the constitutive NADPH oxidase activity elicited by the Trimera and measured with the cytochrome *c* assay by Masoud *et al.* [6]. Addition of PMA to the COSNOX cells expressing only Trimera did not influence the ROS production (data not shown). The addition of DPI, a NADPH oxidase inhibitor, at the end of the measurement caused an immediate and steep drop in the luminescence signal showing an efficient inhibition of the oxidase activity (Figure S3).

**Figure 2.**
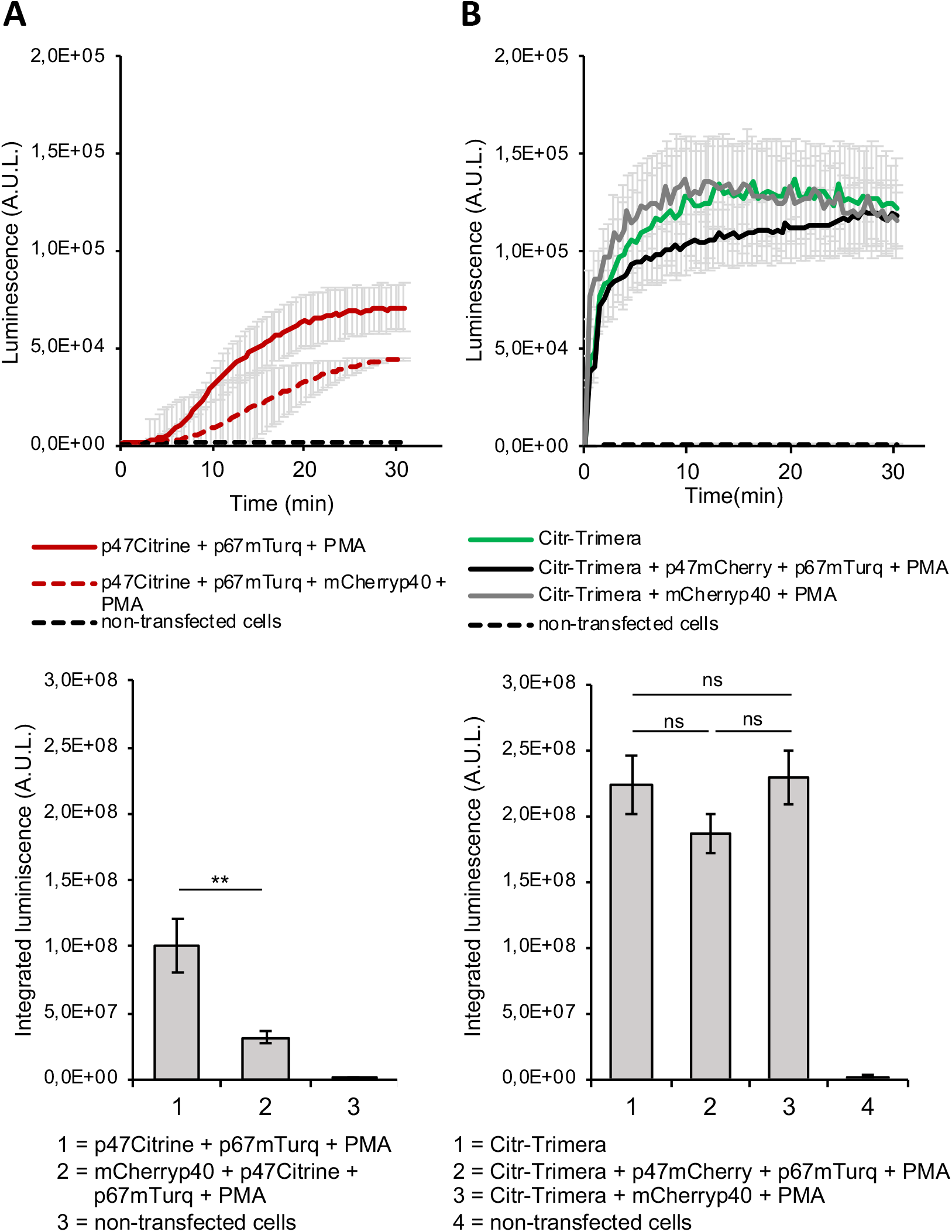
**A:** Top: Effect of p40^phox^ on the ROS production in COSNOX cells transfected with individual NOX subunits. Representative luminometry experiment (duplicates, error bars show SD). Activation by 1 μM PMA. Bottom: Integrated luminescence signal after 30 min of measurement. Error bars show SEM for 4 independent experiments. Statistical analysis performed by t-test (** means p < 0.01). **B:** Top: Effect of the individual NADPH oxidase subunits on the Trimera-elicited ROS production in COSNOX cells. Representative luminometry experiment (duplicates, error bars show SD). Activation by 1 μM PMA. Non-transfected COSNOX cells were used as a negative control. Bottom: Integrated luminescence signal (total ROS production) after 30 min of measurement. Error bars show SEM for 3 independent experiments. Statistical analysis performed by one-way ANOVA.

In order to evaluate if the separate cytosolic subunits may influence the constitutive activity of the Trimera-induced NADPH oxidase, either p47^phox^ and p67^phox^ or p40^phox^ were co-expressed with the Trimera in COSNOX cells (Figure 2B). After 15 min of activation by PMA, the ROS production levels were very similar to the level reached by cells expressing only the Trimera. Comparison of the total ROS production in a 30 min measurement showed no significant difference between these conditions (Figure 2B, *bottom*). mCherryp40^phox^ alone has no effect on the Trimera ROS production. For the separated cytosolic subunits, p47^phox^mCherry and p67^phox^mTurquoise, activated by PMA, we can assume that once the Trimera is expressed in cells, it occupies most of gp91^phox^/p22^phox^ “docking sites” in the membrane, hindering the formation of additional active complexes. The Trimera may also anchor in the same membrane sub-compartment than gp91^phox^/p22^phox^ and “cover” their cytosolic side leading to a steric hindrance and limiting access to p47^phox^/p67^phox^ to p91^phox^/p22^phox^ subunits. So, we were wondering if the Trimera requires the presence of gp91^phox^ or p22^phox^ to locate at the plasma membrane.

### Trimera localizes in the plasma membrane independently of gp91^phox^ and p22^phox^

We already observed the location of the Trimera at the plasma membrane of COSNOX cells [6]. We now compare the localization of the fusion protein Citrine-Trimera in three cell lines: COS-7, COS^p22^ and COSNOX (Figure 3); COS-7 cells do not contain any of the NADPH oxidase subunits except Rac, COS^p22^ cells stably express the p22^phox^ subunit and COSNOX were described in previous paragraph. Using confocal microscopy, we observed that the localization in the three cell lines was identical: the Citrine-Trimera localized predominantly at the plasma membrane. The fluorescent aggregates found in cytosol and nucleus may be due to the binding of non-prenylated Rac1 to the nucleocytoplasmic shuttling protein SmgGDS protein as it was already observed [12], [13]. These results suggest that the presence of neither gp91^phox^ nor p22^phox^ is necessary for the Trimera binding to the membrane.

**Figure 3.**
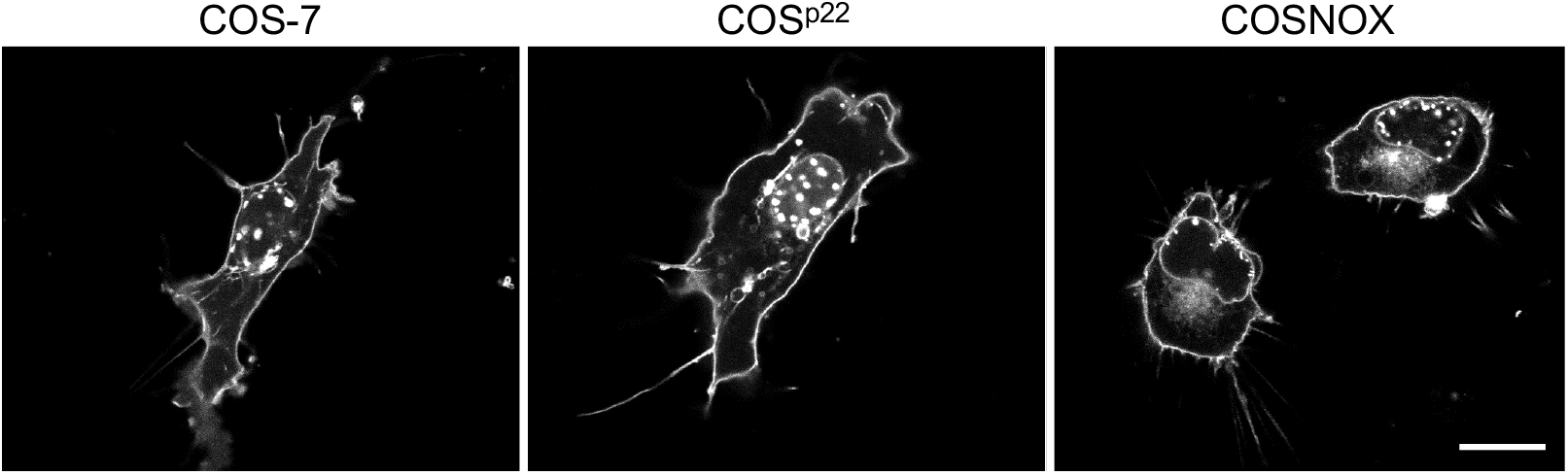
Trimera localization in cells. Confocal images of Citrine-Trimera in COS-7, COS^p22^ and in COSNOX cells. Same scale bar for the three images, 10 μm.

### Continuous NADPH oxidase activity causes an overall decrease of intracellular pH

When Trimera protein is expressed in COSNOX cells, it binds to the plasma membrane, joining the two membrane subunits and turning the NADPH oxidase into the active state. The continuous NADPH oxidase activity leads not only to the O_2_^•−^ overproduction outside of the cell but also to a massive release of H^+^ in the cytosol upon NADPH oxidation into NADP^+^. In this paragraph, we focus on the pH variations induced by the continuous NADPH oxidase activity.

We measured the intracellular pH in COSNOX cells with the ratiometric fluorescent SNARF-1 probe, whose fluorescence is pH sensitive, under different conditions using flow cytometry. In these experiments, the Trimera was labelled with mTurquoise (called mTurq-Trimera), a FP whose fluorescence is compatible with the concomitant observation of pH with SNARF-1. We took the non-transfected COSNOX as the reference with an average pH value of 7.2 (Figure 4A). Control cells expressing only mTurquoise alkalinized slightly, but not in a significant way. pH decreased significantly by 0.2 pH unit in cells expressing mTurq-Trimera. When we added DPI, to the cells expressing mTurq-Trimera, we observed a restoration of the initial pH already after 15 min. This restoration is complete after 45 min (Figure 4A). We controlled that we did not observe any pH recovery in cells expressing mTurq-Trimera treated with DMSO, the DPI solvent. For the negative control, we expressed mTurq or mTurq-Trimera in COS-7 cells and we did not detect any decrease of the intracellular pH (Figure 4B). These findings confirm that NADPH oxidase activity is indeed the source of the cytosol acidification.

**Figure 4.**
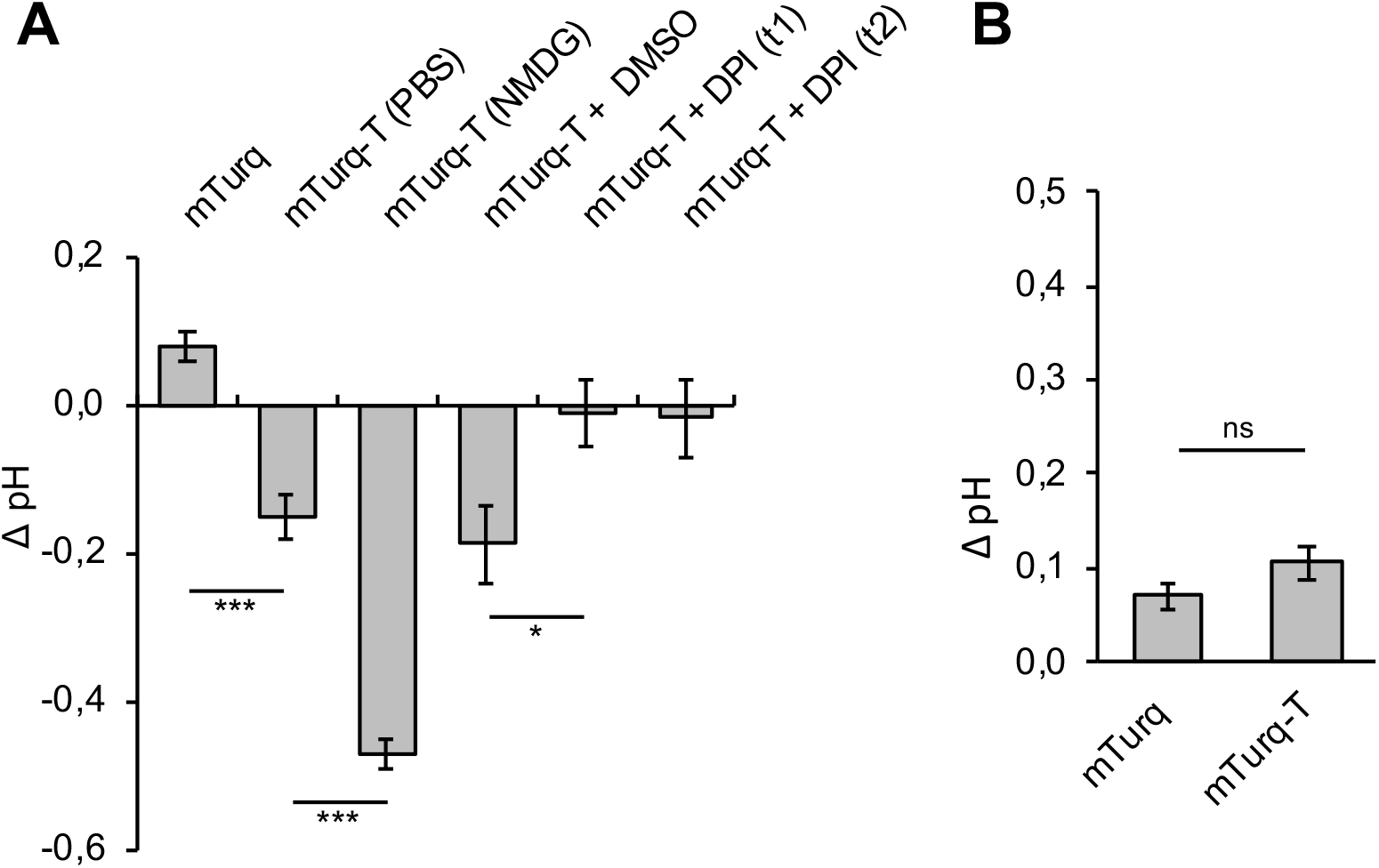
Continuous NADPH oxidase activity causes acidification of the intracellular pH in COSNOX cells. **A:** Comparison of ΔpH in COSNOX cells 24 h after the transfection by mTurquoise-Trimera or mTurquoise. ΔpH was calculated always against the intracellular pH of non-transfected COSNOX cells (average pH value of 7.16). The DPI was added 15 min (t1) and 45 min (t2) before pH measurement. Means and errors bars (SEM) for 5 independent experiments. Statistical analysis performed by t-test (***p<0.001). **B:** Negative control. Comparison of ΔpH in COS-7 cells 24 h after the transfection mTurquoise-Trimera or mTurquoise. Errors bars show SEM for 3 independent experiments. Statistical analysis performed by t-test.

The observed acidification raised the question of the pH regulation in the COS-7 cells used in the study. Indeed, on the contrary to professional phagocytes, in which the pH variations due to the NADPH oxidase activity are counter balanced by proton channels, the COS-7 cells do not have such channels [14]. Nevertheless, COS-7 cells express a Na^+^/H^+^ exchanger and we explored its potential role in the cellular pH equilibration. These experiments required a buffer devoid of Na^+^, to avoid any contribution of the Na^+^/H^+^ exchanger to the pH equilibration. The cells expressing mTurquoise-Trimera were thus incubated a N-Methyl-D-glucamine buffer (NMDG buffer, see Material and Methods) for 1h just before the pH measurement. As shown in Figure 4A cells in NMDG buffer acidified markedly, with their cytosolic pH decreasing by 0.5 pH unit. These data indicate that the Na^+^/H^+^ exchanger extrudes the H^+^ generated in the cytosol by NADPH oxidase and thus is necessary for equilibrating pH fluctuations.

### Continuous NADPH oxidase activity triggers apoptosis

Furthermore, the continuous activation of the NADPH oxidase leads to a massive O_2_^•−^ overproduction. Part of the cocktail of ROS produced, may cause oxidative stress, damage proteins, nucleic acids, lipids, membranes and organelles and finally affect cell survival [15].

We first evaluate the influence of this massive production on cell viability. These experiments have been performed using Annexin V as apoptosis marker and the cells were analyzed by flow cytometry (Figure 5). We investigated the percentage of apoptotic cells within the COSNOX cell population expressing Citrine-Trimera 24 h after transfection. As negative controls, we used COSNOX cells expressing Citrine or COS-7 expressing Citrine-Trimera. To compare highly fluorescent cells in the Citrine channel with cells showing lower fluorescence intensities, we defined two zones in the quadrant Q2 of the dot plots, Q2’ and Q2’’ The percentage of apoptotic cells (quadrant Q2) among Citrine positive cells (quadrant Q2 + Q3) were calculated from the dot plots. The percentage of apoptotic cells was significantly increased in the cells expressing Citrine-Trimera compared to cells expressing Citrine only (Figure 5A and Figure 5B, *left*). In COS-7 cells that do not contain any of the membrane subunits of the NADPH oxidase, very few apoptotic cells were detected in Q2’, whereas a small population of apoptotic cells was detected in Q2’’, the zone of high Citrine-Trimera expression (Figure 5A, *right*). By contrast, in COSNOX cells a significantly increased level of apoptotic cells was observed in Q2’ showing the effect of ROS produced by active NADPH oxidase (Figure 5B, *right*). The percentage of apoptosis reached even higher levels in the Q2’’ zone of COSNOX cells expressing Citrine-Trimera as compared to COS-7 cells (Figure 5B, *right*). Taken together, these results indicate that long-term continuous NADPH oxidase activity and subsequent oxidative stress induces apoptosis in cells.

**Figure 5.**
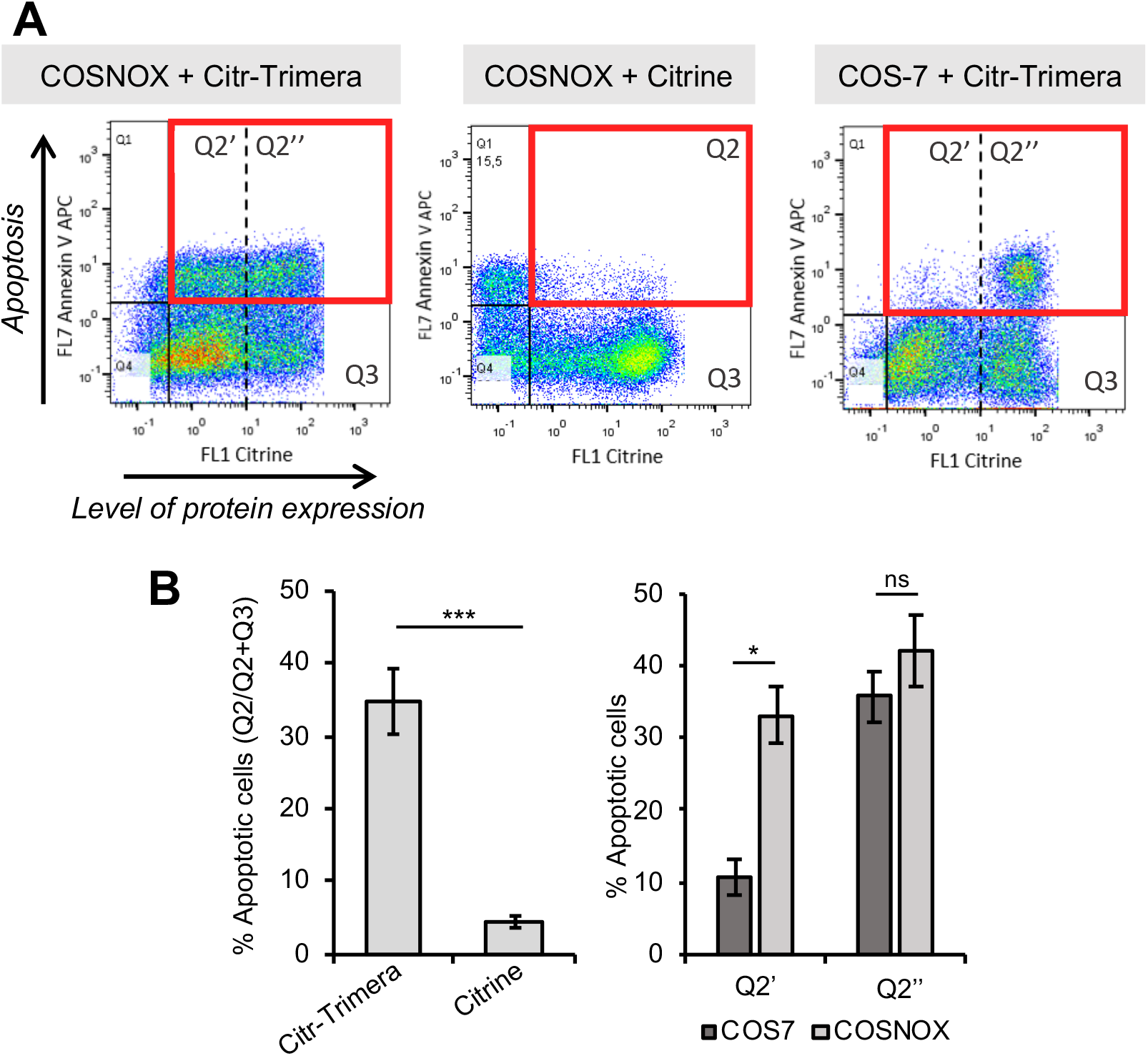
Influence of the continuous NADPH oxidase activity on cell viability. **A:** Apoptosis in COSNOX/COS-7 cells 24 h after the transfection with Citrine-Trimera or Citrine. The dot plots show representative experimental data. The quadrant Q2 (in red) contains cells expressing the protein of interest and being apoptotic as well. Q2’ contains cells with moderate expression level of Citrine-Trimera. Q2’’ contains cells with high expression level of Citrine-Trimera. **B:** Left: Mean % of apoptotic cells in Q2 in COSNOX cells. Right: Comparison of mean % of apoptotic cells in Q2’ and Q2’’ for COSNOX or COS-7 cells transfected with Citrine-Trimera. Errors bars show SEM for 4 independent experiments. Statistical analysis performed by t-test and one-way ANOVA test followed by a Tukey’s Multiple Comparison Test (*** means p < 0.001, * means p < 0.05).

### Increased levels of ROS induce lipid peroxidation of plasma membrane

In this next step, we were wondering if the elevated levels of oxidative stress could induce lipid peroxidation of the plasma membrane. Indeed, the continuous NADPH oxidase activity elicited by the Trimera located at the plasma membrane decreased the cell viability by causing apoptosis that could be triggered by lipid peroxidation [16]. The peroxidation may alter lipid asymmetry and lipid polarity, which affects membrane surface, membrane proteins and enzymes [17]. As a consequence, changes in the membrane thickness, membrane fluidity, and membrane permeability may appear and disrupt normal cell metabolism [18].

First, we opted for an immunostaining strategy to detect 4-hydroxynonenal (4-HNE), one of the products of lipid peroxidation, which is commonly detected as a marker of the process [16, 19, 20]. Second, we used a super-resolution infrared (IR) technique derived from an atomic force microscopy (AFM), called AFM-IR [21] in order to evaluate, at the nanometer scale, the morphological and chemical modifications that occur in the plasma membrane.

To detect 4-HNE, we used a primary antibody anti-4-HNE in combination with a secondary IgG antibody labeled by Alexa Fluor 647. Fluorescence intensity of Alexa Fluor 647 (Figure S4), which corresponded to the amount of 4-HNE in cells, was analyzed by confocal microscopy. The positive control - COSNOX cells treated by the external addition of H_2_O_2_ showed significantly increased fluorescence intensity in comparison to all other conditions. However, there was no significant difference in 4-HNE levels between COSNOX cells transfected with Citrine-Trimera or Citrine alone compared to the non-transfected cells. Thus, the continuous NADPH oxidase activity does not engender distinctly increased levels of 4-HNE in COSNOX cells.

The second approach to search for morphological and chemical modifications in the plasma membrane induced by the continuous NADPH oxidase activity was the nanospectroscopy AFM-IR (Figure 6). This method is a hybrid technique combining high spatial resolution of an AFM (down to 20 nm) depending on the diameter probe (tip) and the chemical analysis of IR spectroscopy (see detail in Material and Methods and Figure S6A). Here, the chemical composition of the sample is detected at the same resolution as the AFM images. This method allows to acquire either AFM images (topography of the surface – morphological analysis), IR chemical maps at a specific wavenumber (location of specific compounds within the cell) or very local IR absorption spectra (to determine the chemical composition and/or the effect of the micro-environment) at a chosen region of interest (ROI) of the sample (Figure 6B). Biomolecules, such as nucleic acids, proteins, lipids, and carbohydrates can be directly probed without any labeling as they possess their own typical vibrational fingerprints in the mid-IR region (Figure S6B). After lipid peroxidation, a huge variety of products like alcohols, ketones, aldehydes, and ethers might be found [22]. Two new chemical features, that might be observed, are the appearance of hydroperoxy (C-O-OH) or carbonyl (C=O) groups in diverse oxidized phospholipid products. The IR absorption of the C=O bond is extremely sensitive to its environment and to the other chemical groups present in its vicinity. In the literature, a band in the region 1755–1720 cm^−1^ is generally assigned to the C=O bond in functional groups like esters, aldehydes, ketones and is usually described as a signature of lipid peroxidation in various biological samples [23–25].

**Figure 6.**
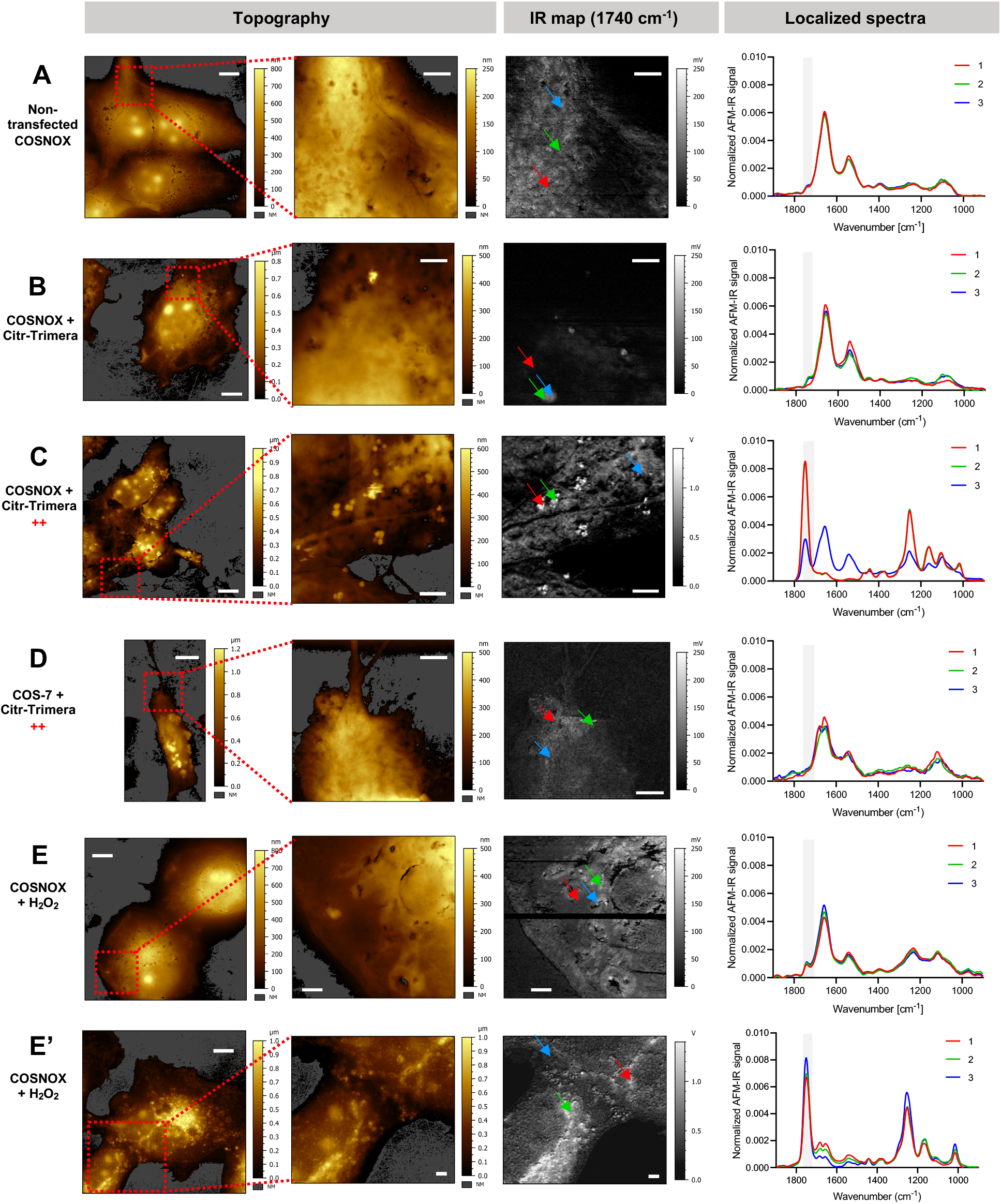
Detection of chemical modification caused by lipid peroxidation in cell plasma membrane by AFM-IR. ‘Topography’ column shows the topography of a large region containing the analyzed cells and of a zoomed selected area (red dashed square). The pseudocolor scale reflects the height of the cell. Column ‘IR map (1740 cm^−1^)’ shows the IR map of the zoomed region. A: Non-transfected and non-treated COSNOX cells. B: COSNOX cells transfected with a normal amount of the Citrine-Trimera DNA. C: COSNOX cells transfected with a high amount of the Citrine-Trimera DNA (three times more than for normal transfection as in case of B). D: COS7 cells transfected with a high amount of the Citrine-Trimera DNA. E + E’: COSNOX cells treated with 200 μM H_2_O_2_ overnight. The last column shows three representative spectra for each condition. Arrows (colored with corresponding colors) in the zoomed IR map show acquisition sites of these spectra. Spectra were normalized to the area under the curve. The region concerning the lipid peroxidation-related signal (1740-1755 cm^−1^) is highlighted in grey. The data and the images were processed using Orange and Mountain map software respectively. Scale bars 10 μm for large topographies, 2.5 μm for zoomed regions.

We investigated COSNOX and COS-7 cells expressing Citrine-Trimera. We incubated cells with H_2_O_2_ as a positive control (Figure 6E, 6E’), whereas non-transfected cells were the negative one (Figure 6A). First, the fluorescent cells expressing Citrine-Trimera or Citrine were identified by wide-field fluorescence microscopy and then analyzed by AFM-IR. For each condition, we acquired a topography showing the surface of the cells (Figure 6, *left columns*). Then a zoomed region of interest (100 to 200 μm^2^) in the membrane, away from the nucleus, was chosen to collect chemical information (IR absorption of the membrane). An IR map at a specific wavenumber was recorded for the whole zoomed ROI (Figure 6, *middle right*). At a chosen location (specified in Figure 6), within the zoomed ROI, full AFM-IR (local IR measurements) spectra from 1900 to 900 cm^−1^ were acquired (Figure 6, *right*).

The local IR spectra for the non-transfected COSNOX cells present the strong characteristic IR bands of the protein peptide bond (amide I at 1660 cm^−1^ and amide II at 1540 cm^−1^) and a low absorption in the fingerprint region corresponding mainly to other bonds in proteins and lipids (C-H bending at 1480-1350 cm^−1^) and at lower wavenumber characteristic bands for carbohydrates and phosphates (Figure 6). We transfected the COSNOX cells with two different quantities of plasmid DNA to induce different expression levels of the Citrine-Trimera (noted Citrine-Trimera and Citr-Trimera++ in Figure 6B and Figure 6C respectively). A pattern very similar to non-transfected cells is observed for COSNOX cells expressing low level of Citrine-Trimera (Figure 6). Nevertheless, at some localization, a slight band at 1740 cm^−1^ appeared associated with a modest increase of absorption in the low fingerprint region (Figure 6, *2 and 3*). These features are much more pronounced when the level of Trimera increased (Figure 6). The strong band at 1740 cm^−1^ is associated with four well defined bands in the fingerprint region at 1260-1250 cm^−1^, 1171-1160 cm^−1^, 1110-1100 cm^−1^ and 1020-1010 cm^−1^. Especially, the strong absorption band observed at 1260 cm^−1^ could be assigned to the C-O stretching (mixed with the O-H in plane bending) of lipid hydroperoxides [17]. For the positive control, COSNOX cells treated overnight with 200 μM of H_2_O_2_, we observed some heterogeneity between cells. In some, a moderate signal at 1740 cm^−1^ was detected (Figure 6E), whereas in others a strong signal at 1740 cm^−1^ associated with the new bands in the fingerprint region was observed (Figure 6E’). The negative control, COS-7 cells transfected with high amount of Citrine-Trimera showed very similar IR fingerprint with almost non-detectable peak at 1740 cm^−1^ as non-transfected COSNOX cells (Figure 6D).

The IR maps recorded at 1740 cm^−1^ revealed the location of the peroxided species (Figure 6, *middle right*). In non-transfected COSNOX cells and COS-7 cells transfected with Citrine-Trimera, a homogeneous low signal on the 1740 cm^−1^ IR map was consistent with the spectra (Figure 6A, Figure 6D). The white hotspots in COSNOX cells expressing Citrine-Trimera corresponded to strong and localized absorption at 1740 cm^−1^ (Figure 6). The amount and the intensity of hotspots increased for the highest Trimera expression level. For control cells exposed to H_2_O_2_, the absorption at 1740 cm^−1^ seemed to be more homogenously distributed with smaller and diffuse hotspots compared to COSNOX cells expressing a high amount of Trimera (Figure 6E’).

The local spectra indicated that the IR bands at 1740 cm^−1^ as well as at 1260-1250 cm^−1^, 1171-1160 cm^−1^, 1110-1100 cm^−1^ and 1020-1010 cm^−1^ were also present in the cells exposed to H_2_O_2_; this band being likely due to the lipid peroxidation as described in the literature [23]. The relative intensity of those bands is strongly linked to the final mixture of peroxides that may slightly differ from the one obtained in COSNOX cells expressing Citrine-Trimera. The negative controls confirmed that the peak at 1740 cm^−1^ was directly related to the expression level of Citrine-Trimera in the COSNOX cells. The comparison of different expression levels of the Trimera protein suggested that the presence of those bands was most probably due to lipid peroxidation caused by intensive NADPH oxidase activity.

This chemical signature of oxidative stress is less specific than the detection of 4-HNE. The immunostaining of 4-HNE detects a single oxidized species, whereas the IR signal at 1740 cm^−1^ may correspond to many different oxidized molecules. The IR signal likely reflects the molecular diversity of the chemical modifications induced by the sustained NADPH oxidase activity. An interesting finding was the localized distribution of the oxidation in hotspots of a few hundreds of nm. This was quite surprising as we rather expected a homogenous ROS production in the cell membrane.

### Trimera forms clusters in the plasma membrane independently of gp91^phox^

In order to understand the localized oxidative damages observed with the AFM-IR technique, we investigated the Trimera membrane distribution. To this aim, we decided to exploit the Förster Resonance Energy Transfer (FRET) between two fluorophores, a donor, mTurquoise, and an acceptor, mCitrine fused to the Trimera and expressed together (Figure 7A). The fluorescence emission spectrum of mTurquoise overlays the absorption spectrum of mCitrine, so both FPs fulfill the spectral condition for FRET: upon their excitation, the donors, in the excited state, can transfer part of their energy to the acceptors in the ground state, which in turn become fluorescent. The FRET efficiency is highly dependent on geometrical parameters such as the donor/acceptor distance, which must be less than 10 nm (Figure 7A) and on protein topology. Consequently, the FRET efficiency between donors and acceptors fused to the protein of interest and co-expressed in the cells can be used to evaluate the distribution of those proteins at the membranes, where the average spacing depends on their concentration and lateral distribution.

**Figure 7.**
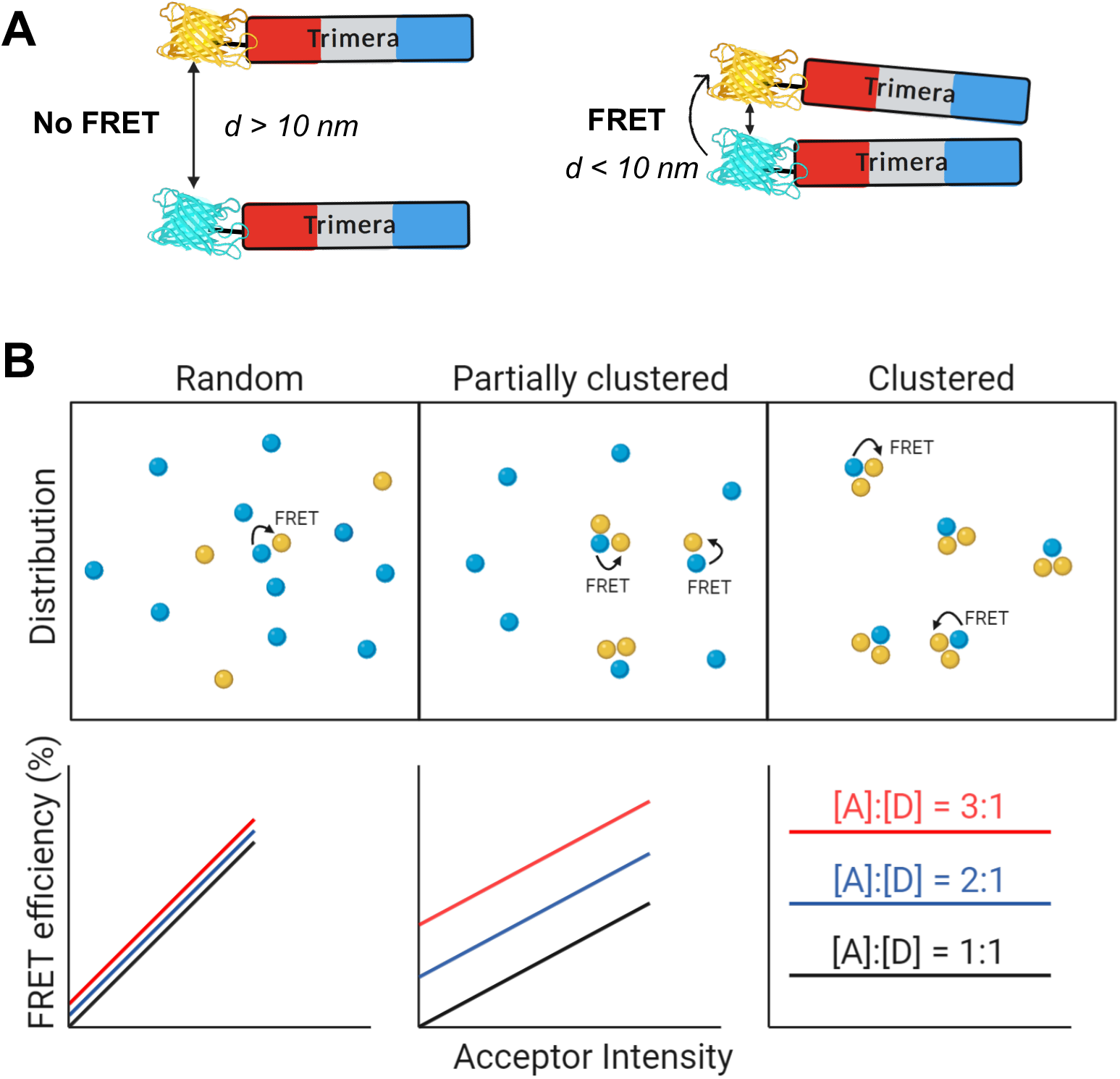
Investigation of protein distribution in membrane of living cells by FRET. **A:** In case that Trimera proteins labeled by a donor (mTurquoise) or by an acceptor (mCitrine) are in close proximity, the FRET phenomenon will occur. **B:** Possible distributions of membrane-bound proteins (top) and associated FRET efficiency curves (bottom). Adapted from [28]. In our study, we measured the FRET variations by FLIM.

Proteins can be homogeneously and randomly distributed in the membrane, they can form clusters, or they can create a mixed distribution containing clusters and free molecules (Figure 7B, *upper row*). The organization at the molecular level influences the possibility of FRET to occur: an efficient FRET can be observed if two molecules are close either due to clustering, or due to high concentrations and packing, which bring them close enough even in the absence of microdomains [26], [27]. Consequently, the FRET efficiency varies with the density of labelled proteins in the cell membrane [28].

In a random distribution, the FRET efficiency increases linearly with the acceptor surface density that is proportional to its fluorescence intensity. By contrast, in case of a clustered distribution, the FRET efficiency will depend on the relative quantity of donors and acceptors in the cluster. If the clusters are small enough or sufficiently diluted, there will be no energy transfer between them. Thus, the FRET efficiency will be independent of the acceptor intensity for given acceptor-to-donor ratios. The intermediate situation concerns a partially clustered distribution. So, monitoring FRET efficiencies as a function of the acceptor fluorescence intensity for different acceptor-to-donor ratios reveals a pattern characteristic for each type of distribution curve (Figure 7B, *bottom row*) [28].

First, we validated our experimental approach on a model system using myristoylated and palmitoylated (MyrPalm) fluorescent proteins, which are known to clusterize in the plasma membranes [26]. We transfected the COS-7 cells simultaneously by MyrPalm-Aquamarine (donor) and MyrPalm-YFPA206K (acceptor). The DNA concentration of the MyrPalm-YFPA206K used for transfections was modulated in order to expand the range of acceptor fluorescence intensities and various [A]/[D] ratios. For each cell, the donor fluorescence lifetimes and donor and acceptor fluorescence intensities were measured successively. The apparent FRET efficiency (*E_app_*) was evaluated from the donor lifetimes using Equation 3 (see MM) and the relative acceptor-to-donor concentration ([A]/[D]) was deduced form the intensities thanks to the custom calibration (see SI). Cells were classified in three categories depending on the [A]/[D] ratio: 0-1, 1-3 and > 3.

Our results show different levels of FRET efficiency for each relative concentration [A]/[D] ratio. Those efficiencies are predominantly independent of the acceptor fluorescence intensity, the slopes of the fitting curves being very low (Figure 8A, *left*). To modulate the distribution of MyrPalm proteins in the membrane, the cells were treated with 10 mM 5-methyl-β-cyclodextrin (MβCD), a treatment known to disrupt lipid rafts and caveolae by depletion of cholesterol [29]. The slopes of fitting curves drastically increased, they became three times larger (Figure 8C, *left*), indicating the alteration of protein distribution and a decrease in the level of clustering as expected for MyrPalm-FPs proteins that are known to cluster in lipid rafts. Thus, our experimental workflow was suitable for the investigation of protein clustering.

**Figure 8.**
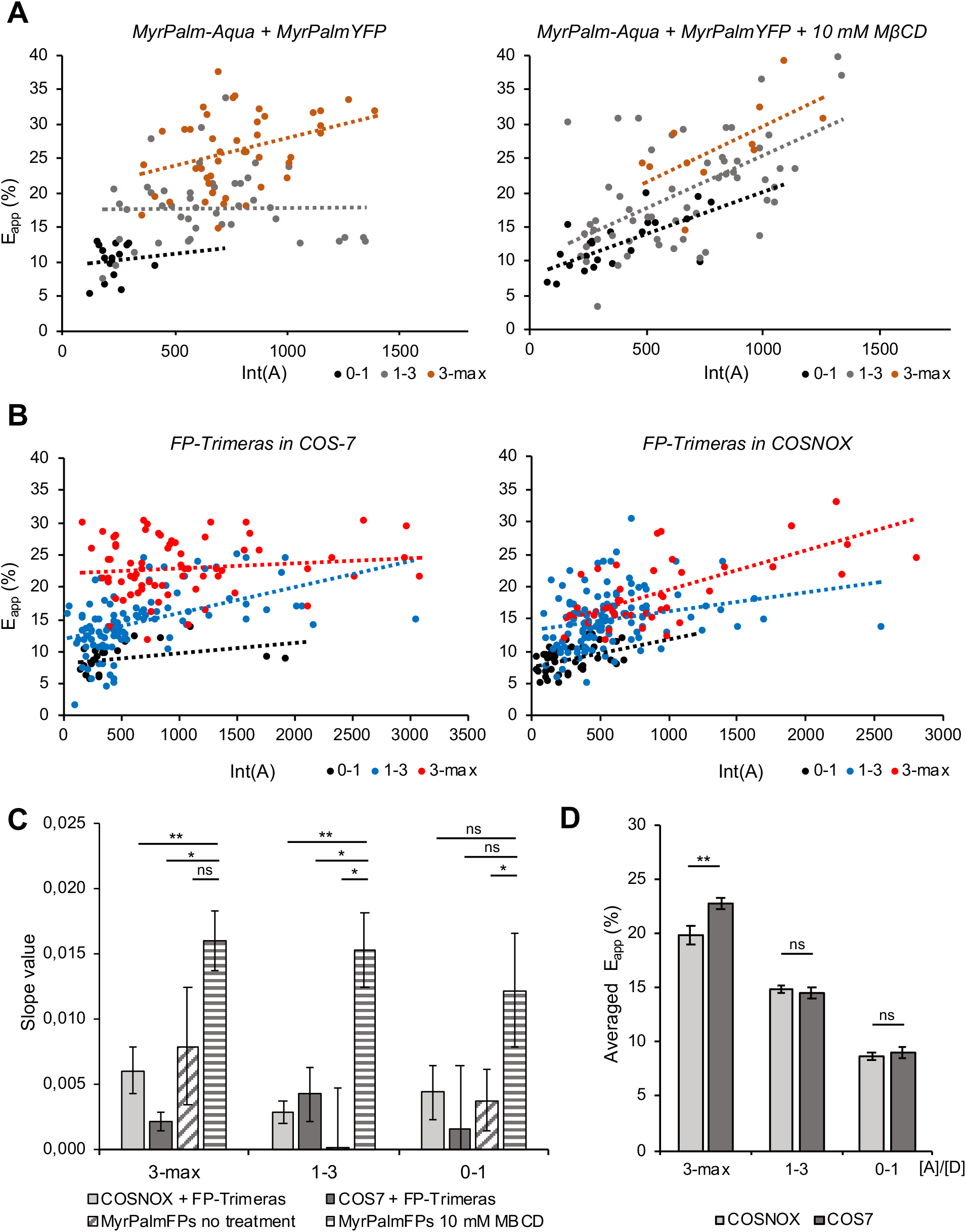
Distribution of MyrPalm proteins (model system) and Trimera proteins in living cells monitored by FRET-FLIM. Graphs show FRET efficiency (E_app_) in function of the acceptor intensity. Cells were classed in three different [A]/[D] ratios (different colors). **A:** COSNOX cells transfected by MyrPalm-Aquamarine and MyrPalm-YFPA206K were analyzed without any special treatment (left) or they were treated with 10 mM MβCD (right). **B:** mCitrine-Trimera and mTurquoise-Trimera were expressed in COS-7 cells (left) or in COSNOX cells (right). **C:** Comparison of slopes of fitting curves for FP-Trimeras and MyrPalm-FPs FRET-FLIM experiments. **D:** Comparison of averaged E_app_ between COSNOX and COS-7 cells for selected range of the acceptor intensity: Int(A) = 500-900. Error bars show SEM for n = 3 experiments. Statistical analysis performed by one-way ANOVA followed by a Tukey’s Multiple Comparison Test (** means p < 0.01).

Therefore, we used the same analytical workflow to investigate the FP-labeled Trimera distribution in the plasma membrane. mTurquoise-Trimera and mCitrine-Trimera were expressed in COS-7 or COSNOX cells in order to evaluate a possible implication of the membrane subunits (gp91^phox^ and p22^phox^) in Trimera spatial organization at a subcellular level. Different ratios of [A]/[D] were generated as previously for MyrPalm-FPs. During the data treatment, the cells were classified into three categories depending on the [A]/[D] ratio. The *E_app_* for each category was plotted against the acceptor fluorescence intensity as shown in Figure 8B.

The slopes of the fitting curves obtained for both the FP-Trimeras in COS-7 or COSNOX cells were close to those of non-treated MyrPalmFP proteins in each [A]/[D] category (Figure 8C, *left*). These findings suggest a mainly clustered distribution of the FP-Trimera in both cell lines.

To assess if COS-7 and COSNOX cell lines show genuinely the same distribution in this experiment, averaged *E_app_* for every [A]/[D] ratio (Figure 8C, *right*) was compared. It increased systematically with the growing [A]/[D] ratio and the highest *E_app_* values (~ 20%) were found for the category 3-max of [A]/[D]. This is also seen via the relative position of the linear fits of the experimental dots: the red fitting line for the highest ratio is above the blue and the black one. This is likely a consequence of the increase of the intra-cluster FRET efficiency with the [A]/[D] ratio as it is illustrated in Figure 7B (arrows): statistically, within a cluster, the number of potential FRET acceptors increases in the vicinity of each donor. For the category 3-max of the [A]/[D] ratio, we observed a significant difference in the average *E_app_* between COS-7 and COSNOX; average *E_app_* values being higher for COS-7 cells (Figure 8D). Since we detected the ROS production in COSNOX cells expressing Trimera, the latter has to interact directly with gp91^phox^/p22^phox^ complexes. We can speculate that the presence of gp91^phox^/p22^phox^ in the Trimera clusters in COSNOX cells may impose topological constraints on the relative A/D distance for example. These constraints have certainly consequences on the intra-cluster FRET efficiency, as they could be observed only for the highest [A]/[D] ratio (Figure 8C, *right*), when the contribution of the intra-cluster FRET is likely predominant. The specificity of the Trimera-gp91^phox^ interaction requires further experimental investigations, which might also help to explain the role of gp91^phox^ in the clustering process.

## DISCUSSION

### Membrane targeting and constitutive ROS production

We showed that Trimera localizes to the plasma membrane independently of gp91^phox^/p22^phox^ subunits and this result raises the question of the membrane targeting of the Trimera. The membrane targeting of the small G-protein Rac1 is ensured by the exposition of its polybasic region (PBR) in a GTP-bound form and by the attachment of a lipid anchor to its C-terminus [30]. In the Trimera sequence, Rac1Q61L is present at the C-terminus (Figure 1). The mutation Q61L ensures that Rac1 is permanently in a GTP-bound, constitutively active form and prenylated by a geranylgeranyl chain. Within this frame, we could suggest that Rac1Q61L is the principal driving force in the Trimera translocation to the plasma membrane. In addition, the PX domain of p47^phox^ also present in the Trimera can interact with phosphatidylinositol (3,4)-bisphosphate (PI(3,4)P_2_) in the membrane [31].

In COSNOX cells, Trimera elicits a continuous activity of the NADPH oxidase. It means that Trimera adopts a position allowing the activation domain of p67^phox^ to interact with gp91^phox^ and resulting in ROS production. In the following, we discuss the adequacy of the Trimera structure to trigger the observed continuous oxidase activity. During the NADPH oxidase activation process, the TPR motifs of p67^phox^ bind to Rac1-GTP. The Rac1-GTP-p67^phox^ complex facilitates interaction of gp91^phox^ with p67^phox^ via its activation domain [32]. In Trimera, Rac1-GTP (active form due to the Q61L mutation) and 1-212 aa residues of p67^phox^ are directly fused without any linker (Figure 1), making the activation domain of p67^phox^ ready to interact with gp91^phox^, if they co-localize in the membrane and have the right orientation. It has been shown previously that p47^phox^ can also directly interact with gp91^phox^. Regions in p47^phox^ involved in that interaction have been identified only at one site situated within the AIR domain (aa 323–342) [33]. In Trimera, only aa residues 1-286 of p47^phox^ are present (from the PX domain to the C-terminus of the second SH3 domain). So, the region that has affinity for gp91^phox^ is missing. However, although p47^phox^ does not directly interact with gp91^phox^ in COSNOX cells, it could still interact with p22^phox^ by its SH3 domains that are conserved in Trimera protein. The p47^phox^-p22^phox^ interaction is an essential interaction that can help to find the right orientation of Trimera to dock gp91^phox^ and enable ROS production. Moreover, p47^phox^ targets PI(3,4)P_2_ [34] in the cell membrane and Rac1 targets anionic phospholipids including PI(3,4)P_2_. This increases the chance for optimal Trimera-gp91^phox^ interaction. Importantly, the co-localization of gp91^phox^ and Trimera in the plasma membrane is mandatory to initiate the NADPH oxidase activity.

### Constitutive ROS production: how much is too much?

While low H_2_O_2_ levels (1–10 nM) play an important role in redox signaling, higher levels (> 100 nM) overwhelm the anti-oxidant defenses causing oxidative stress and may cause irreversible cell damage [35]. Masoud *et al.* measured with the quantitative cytochrome *c* assay that the superoxide anion production rate elicited by Trimera is 11 nmol.min^−1^ for 10^7^ cells [6]. Estimating that the expression elicited by Trimera started 10 h after the transfection itself (real-time observation, data not shown), we can calculate that each cell has produced 0.9 pmol of superoxide anion within 24 h after the transfection. If we hypothesize that the superoxide anions quickly dismutate into H_2_O_2_ and that 0.1 to 1% of H_2_O_2_ diffuse back into the cell, 0.009 pmol correspond to an equivalent intracellular concentration of 0.6 to 6 mM of H_2_O_2_ for a cell volume of 1.5 pL (cell size 30 × 10 × 5 μm). Despite the intracellular antioxidant defense, those values are in the range of a strong oxidative stress and can cause apoptosis in cells [3]. We also observed a small population of apoptotic cells in COS7 cells likely due to the very high overexpression of Trimera at the plasma membrane that may cover the intracellular membrane leaflet and prevent the correct interaction of other membrane bound proteins.

We also reported that the constant NOX activity leads to acidification of the intracellular environment and one might wonder if the pH variation in the cytosol could also contribute to the apoptosis. It has been described that intracellular pH of apoptotic cells is more acid (typically diminished by 0.3-0.4 pH units) than that of healthy cells [36], which is consistent with our experiments. In the two major apoptotic pathways known to date, that is, the ‘mitochondria-dependent’ pathway and the ‘death receptor’ pathway, activation of a caspases has been recognized to play a pivotal role. It was shown that activation of certain caspases is mediated by cytochrome c that is released from mitochondria, when intracellular acidification occurs [37]. Other pro-apoptotic signalling pathways, such as the p38 mitogen-activated protein kinase (MAPK) pathway, might also be affected by a cytosolic acidification [36]. Taken together, we assume that the intracellular acidification may also participate to the initiation of the apoptosis pathway in our experiments. Another event that can contribute to apoptosis is lipid peroxidation.

### Localized oxidations and clustering

Using AFM-IR, we were able to observe the IR signature of chemical modifications at 1740 cm^−1^, attributed to the C=O bond, as well as at 1260-1250 cm^−1^, 1171-1160 cm^−1^, 1110-1100 cm^−1^ and 1020-1010 cm^−1^, which is correlated with the increase of C-O stretching and OH bending of hydroperoxides vibration band. The measurements showed that the intensity of IR signal is directly related to the presence of the Trimera protein and correlated to its expression level. This suggests that the appearance of these bands is due to the intense NADPH oxidase activity. Reactive oxygen species can react not only with lipids, but also with DNA, RNA and proteins. The spectra were recorded by placing the AFM tip on the membrane far from the nucleus, thus oxidation products from DNA and RNA could be excluded. Furthermore, proteins are intricate as oxidation targets, considering 20 different side chains plus the backbone. There are many oxidation pathways that can occur leading to high complexity of products that are formed [38]. On the other hand, lipids have a more limited number of oxidation products and they can massively contribute to the signal at 1740 cm^−1^ [23–25, 39]. We cannot exclude the participation to the IR signal of oxidized forms of proteins, but the strong contribution of lipids from the membrane is expected to be predominant. Interestingly, the signal at 1740 cm^−1^ was detected in organized hotspots with diameter ~ 200 nm. This finding is coherent with our result showing Trimera clustering in the plasma membrane observed by FRET-FLIM. Recently, using *d*STORM super-resolution microscopy Joly *et al.* not only showed that the catalytic subunits (gp91^phox^) also clusterize in the plasma membrane, but they were able to determine the size of the clusters around 60 nm [40].

Technically, notions of clustered distribution of the NADPH oxidase were documented by immunoelectron microscopy already in 1997 [41]. Nowadays, clustering phenomenon seems to be involved in cell signaling hubs and pathways. Co-clustering of the Trimera with gp91^phox^ probably leads to a localized and very intense ROS production leading to the IR signal patches. To evaluate the size of Trimera clusters and understand their role in the formation of the oxidized lipids spots in the plasma membrane, further experiments are needed. Besides, we should also notice that Holman *et al.* investigating lung fibroblasts at different points of their cell cycle and assigned the increase of the IR band at 1740 cm^−1^ to the cells undergoing apoptosis or to already dead cells [42]. This is coherent with our results showing that about 35% of COSNOX cells expressing Trimera become apoptotic.

Our study validated Trimera protein as a suitable tool to simulate the NOX2 active state in living cells. It enables exclusive investigation of the NADPH oxidase in its active phase and can be considered as a potential tool for screening of inhibitors, which are NOX2-isoform specific. Our data suggest that the NADPH oxidase forms clusters and its activity is focused to very local hotspots in the cell membrane, but its upregulation influences overall cell function.

## MATERIALS AND METHODS

### Materials

Horseradish peroxidase (HRP), phorbol 12-myristate 13-acetate (PMA), N-Methyl-D-glucamine (NMDG), nigericin and methyl-beta-cyclodextrin were purchased from Sigma-Aldrich (St Quentin, France). Dulbecco’s PBS (DPBS) and all cell culture media were purchased from Gibco (France). Annexin V APC and propidium iodide (PI) were purchased from Invitrogen (France). Primary antibody anti-4-HNE was purchased from Abcam (mouse, ab48506). Secondary antibody conjugated to Alexa Fluor 647 was purchased from Thermo Fisher Scientific.

### Plasmid library

The plasmid coding for the Citrine-Trimera fusion protein was prepared in our laboratory as described previously [6]. The orginal GFP fused Trimera plasmid was a kind gift of E. Pick [7]. This plasmid was then used for creating mTurquoise-Trimera by exchanging the fluorescent protein tag. Citrine-Trimera and plasmid of mTurquoise (vector, originally pECitrineC1) were digested with *Hind*III and *Kpn*I and ligated using standard protocols. The sequence of the resulting plasmid was verified by sequencing (Genewiz, Germany). Endotoxin free plasmid, which is more convenient for cell transfection, was prepared using the EndoFree Plasmid Maxi Kit (Qiagen, France). For FRET-FLIM experiments, a plasmid of mCitrine-Trimera with monomeric Citrine (A206K mutation) was prepared. Using QuikChange Lightning Multi Site-Directed Mutagenesis Kit (Agilent) and following primers: forward 5’-cctgagctaccagtccaagctgagcaaagacccc-3’ and reverse 5’-ggggtctttgctcagcttggactggtagctcagg-3’, A206K mutation was introduced in the DNA sequence of Citrine.

### Cell culture and transfection

Stock cultures of COS-7 (African Green monkey kidney) and COS^gp91-p22^ cells were maintained at 37°C in DMEM supplemented with 10% fetal bovine serum (FBS) in a humidified atmosphere containing 5% CO_2_.

In the case of COS^gp91-p22^, the culture medium was supplemented with the selecting antibiotics Geneticin (1.8 mg/mL, Thermo Fisher Scientific) and Puromycin (1 μg/mL, Sigma Aldrich). The day before transfection, cells were seeded in selected plates (Greiner Bio-one) and transfected at 80–90% of confluency using X-tremeGENE™ HP DNA Transfection Reagent (Roche) following strictly the manufacturer protocol.

### *In-cell* NADPH oxidase activity assay

The NADPH oxidase activity was measured by luminescence assay. The COS^gp91-p22^ cells were plated in 24-well plates and transfected 24 h before experiment. Transfection efficiency was verified first by wide-field microscopy and then by flow cytometry (Figure S1). Only cells with the same level of transfection were used for the luminescence assay in order to directly compare the ROS production. The assay was performed in PSBG buffer (PBS supplemented with 0.5 mM MgCl_2_, 0.9 mM CaCl_2_, and 7.5 mM glucose) at 35°C using SynergyH1 plate reader (Biotek, USA). L-012 (100 μM, Wako Chemicals) and HRP (20 U/ml) were mixed and added to the wells containing transfected COS^gp91-p22^ cells.

If needed, PMA (1 μM) was added to induces the NADPH oxidase activation process 2 min after addition of L-012 and HRP. 50 μM of DPI was added to stop the O_2_^•−^ production. The ROS production was quantified as integrated luminescence over time.

### pH measurements

A stock solution of SNARF-1 was prepared in DMSO at a concentration of 5 mM and kept at −20 °C.

Cells were seeded in 12-well plates and prior to loading with SNARF-1, the supplemented DMEM was removed and cells were washed with PBS. To load SNARF-1, cells were incubated with 5 μM SNARF-1 for 30 min in PBS at 37°C, 5% CO_2_. Loading was done in PBS buffer to avoid unwanted hydrolysis by esterase present in serum. After incubation, cells were centrifuged (5 min, 1200 rpm), resuspended in PBS and analyzed by flow cytometry. SNARF-1 calibration was performed *in-cell* according to the following method. After incubation with the fluorescent probe, cells were resuspended in buffers containing 15 mM MES, 15 mM HEPES and 140 mM KCl at various pH values (6.0-8.0). The addition of 15 μM nigericin (10 min, RT) allowed an exchange of K^+^ for H^+^ resulting in a rapid equilibration of external and internal pH. The calibration curve varied slightly for each experiment and thus was repeated for each experiment. A typical example is shown in Figure S2.

To exclude any contribution of the Na^+^/H^+^ exchanger, some of the experiments were performed in Na^+^-free medium containing: 140 mM NMDG-Cl, 5 mM KCl, 1 mM MgCl_2_, 1.8 mM CaCl_2_, 10 mM HEPES and 10 mM glucose at pH 7.4 [43]. In this case, cells were first incubated in NMDG buffer for 30 min (37°C, 5% CO2) and then they were loaded by SNARF-1 diluted in the NMDG buffer (30 min, 37°C, 5% CO2). After that, the cells were detached, centrifuged (5 min, 1200 rpm), resuspended in DPBS and analyzed by flow cytometry.

### Apoptosis tests

The cells were seeded in 12-well plates and analyzed 24 h after transfection. Prior to analysis, they were washed with PBS and detached with trypsin (Gibco). For apoptosis tests, cells were resuspended in annexin binding buffer (Invitrogen) containing Ca^2+^ due to the calcium dependence of the Annexin V:PS interaction. Cells were incubated with Annexin V APC for 10 min at RT and analyzed by flow cytometry.

### AFM-IR measurements

COSNOX and COS-7 cells for AFM-IR analysis were grown on the CaF_2_ cover slips in 24-well plates. 24 h hours after transfection (DNA amount: Citrine-Trimera 0,5 μg/well, Citrine-Trimera++ 1,5 μg/well), they were fixed by 4% paraformaldehyde, washed using PBS/water baths to successively decrease the amount of PBS (going from 100% PBS to 100% water) and stored at 4°C before analysis. At least 7 cells per condition were examined using an AFM-IR nano-spectroscopy system. In this technique, the AFM tip is in contact with the sample and detects its thermal expansion induced by the absorption of the IR source. The instrument used in this study is a NanoIR2 (Anasys Instrument, Bruker nano Surfaces, California, USA) combining an AFM set-up in contact mode with an IR pulsed tunable laser covering the mid-IR region from 1945-890 cm^−1^ (QLC beam, MIRcat-QT, DAYLIGHT solutions; peak powers up to 1 W; average power up to 0.5 W, wavelength repeatability < 0.1 cm^−1^ and a tunable repetition rate of 1-2000 kHz). We used a scan rate of 0.5 Hz per line in the contact mode. An Au-coated silicon cantilever (Mikromasch) with a nominal tip radius < 15 nm and a spring constant between 0.03 and 0.2 N/m was used for all measurements (scan rate 0.5 Hz per line, laser power 9%, pulse width 200 ns, pulse rate 766 kHz, PLL (phase locked-loop)). The AFM-IR spectra were collected with a data point spacing of 1 cm^−1^, sweep rate 200 cm^−1^/s, average on 4 spectra at each position (scan rate 0.5 Hz per line, laser power 9%, pulse width 200 ns, pulse rate 766 kHz, PLL) at each wavenumber position over the spectral range 900–1900 cm^−1^.

All experiments were performed at room temperature. The data were analyzed in Orange: baseline correction (linear), gaussian smoothing (SD 5), area normalization; and in Mountain map software: topography and IR-map analysis. Orange is a comprehensive, component-based software suite for machine learning and data mining, developed at Bioinformatics Laboratory, Faculty of Computer and Information Science, University of Ljubljana, Slovenia [44]. Mountain map was developed at Anasys Instruments, USA.

### Immunostaining assay

COSNOX cells were seeded on glass cover slips in a 12-well plate in complete DMEM medium and transfected according to the manufacturer’s protocol. 24 h after transfection, the cells were fixed with 4% paraformaldehyde (Electron Microscopy Sciences) and kept 45 min at 13°C. After washing with PBS buffer, cells were incubated 10 min at RT with 1% glycine (Sigma). Cells were permeabilized by 0.1% saponin (Rectapur Prolabo) diluted in DPBS, 3x 5 min at RT. Afterwards, cells were incubated with DPBS containing 10% FBS and 0.1% saponin for 30 min at RT. The permeabilized cells were incubated with primary antibody anti-4-HNE (Abcam 48506) diluted 1/200 in PBS buffer containing 5% BSA overnight at 4°C. The next day, cells were washed 3x with PBS buffer containing 1% BSA and incubated with the secondary antibody (Alexa Fluor 647, ThermoFisher Scientific) diluted 1/1000 in DPBS buffer containing 1% BSA for 45 min at 37°C. After the final wash with PBS buffer containing 1% BSA, the cover slips were mounted on the glass slides using a mounting medium (ProLong Gold Antifade Reagent).

### Flow cytometry measurements

Transfection efficiency, pH measurements and cell viability were performed with a CyFlow flow cytometer (Partec/Sysmex) equipped with a Flomax software. For the measurement of transfection efficiency, Citrine was excited at 488 nm and detected with a 536/40 nm BP filter, mTurquoise was excited at 405 nm and detected with a 455/50 nm BP filter, and m Cherry was excited at 561 nm and detected with a 610/30 nm BP filter.

For pH measurements, SNARF-1 was excited at 488 nm. The fluorescence of its protonated form was detected with a 590/50 nm BP filter and its deprotonated form with a 675/22 nm BP filter. By calculating a fluorescence intensity ratio of these two forms and using a calibration curve of SNARF-1, final intracellular pH was determined using FlowJo software.

For cell viability, the apoptosis was examined using and Annexin-V APC, which was excited at 635 nm and detected with a 675 nm BP.

### Confocal microscopy

For confocal microscopy imaging, cells were seeded on μ-Slide 8 well glass bottom chamber IBIDI and observed in DPBS. Images were acquired with an inverted Leica TCS SP8X microscope using a 63x oil immersion objective (NA 1.4) and a white laser.

### Fluorescence Lifetime Imaging Microscopy (FLIM)

For FLIM experiments, cells were seeded on glass cover slips (⌀ 25mm) in 6-well plates (Greiner Bio-One) in density 0.2 × 10^6^/mL. They were transfected 24 h prior to experiment. Transfected cells were studied in PBS at 37°C for 2 h maximum in an Attofluor cell chamber (Thermo Fisher Scientific).

Time resolved laser scanning TCSPC microscopy was performed on a custom-made microscope as described previously [45]. The setup is based on a TE2000 microscope with a 60x, 1.2NA water immersion objective (Nikon). The epifluorescence pathway is equipped with an Hg lamp, a set of filter cubes for the different FPs and a CCD camera (ORCA-AG, Hamamatsu Photonics). The TCSPC path is equipped with pulsed laser diodes (440 nm for CFPs, PicoQuant) driven by a PDL800 driver (20 MHz, PicoQuant). The C1 scanning head (Nikon) probes a 100 × 100 μm maximum field of view. To select the FP fluorescence, dichroic mirrors (DM) and filter sets were used before the detection by an MCP-PMT detector (Hamamatsu Photonics). The signals were amplified by a fast pulse pre-amplifier (Phillips Scientific) before reaching the PicoHarp300 TCSPC module (PicoQuant). Counting rates were routinely between 50 000 and 100 000 cts/s.

The lifetime of a fluorophore is independent of its concentration and a precision of a few percent on lifetime is common [46]–[48]. The TCSPC fluorescence decay of all collected pixels in the selected ROIs was computed using the SymPhoTime software (PicoQuant). As donor, we used fluorescent proteins Aquamarine or mTurquoise, whose fluorescence decay can be fitted with a mono-exponential fit function:

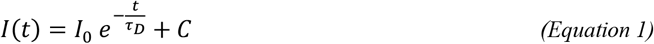

where *I_0_* is the fluorescence intensity at t = 0, τ_D_ is the fluorescence lifetime of a donor alone, and *C* is the constant background.

During the FRET experiment, donor and acceptors are co-expressed in cells. Some donors interact with acceptors, but some may not have any FRET partner. This situation leads to a fluorescence decay fitted with a bi-exponential function:

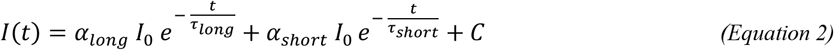

where α_long_ and α_short_ are the proportions of a long (*τ*_long_) and a short (*τ*_short_) lifetime components respectively (SI of [49]).

We calculated < *τ*_DA_ >, the average lifetime of the donor in presence of the acceptor, as:

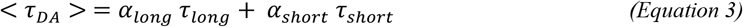

The apparent FRET efficiency (*E_app_*) was calculated using < *τ*_DA_ > and τ_D_:

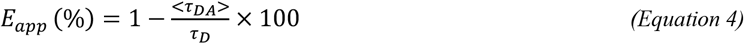

The plotting of *E_app_* against [*A*]/[*D*] required the calibration of fluorescence intensities measured in both donor and acceptor channel in the same ROI used to compute the fluorescence decays. The ratio of fluorescence intensities, *I*(*A*)/*I*(*D*), was transformed in the concentration ratio of fluorescent proteins, [*A*]/[*D*], using a custom calibration procedure of the microscopy setup (see SI). Image treatment was performed in Image J (NIH, USA).

### Statistical Analysis

Data in graphs are represented as means of at least three independent experiments ±SEM. For pH measurements, the significance was tested with Student’s test, for FRET-FLIM experiments with one-way ANOVA. The statistical analysis was performed using R version 3.5.3.

## Abbreviations

aa: amino acid
AFM: Atomic Force Microscopy
DPI: Diphenylene Iodonium
FRET: Förster Resonance Energy Transfer
FLIM: Fluorescence-Lifetime Imaging Microscopy
FP: Fluorescent Protein
HRP: Horseradish Peroxidase
IR: Infra-Red
NADPH: Nicotinamide Adenine Dinucleotide Phosphate reduced form
NMDG: N-Methyl-D-glucamine
PMA: Phorbol 12-Myristate 13-Acetate
ROI: Region of Interest
ROS: Reactive Oxygen Species

## Acknowledgments

COS^gp91-p22^ cells were a kind gift from Mary C. Dinauer (Washington University School of Medicine, USA). The plasmid of the Trimera was a generous gift from E. Pick (Tel Aviv University, Israel). HV was a recipient of a PhD fellowship from Université Paris-Saclay.

