## Supplementary information for "Consequences of the constitutive NOX2 activity in living cells: cytosol acidification, apoptosis, and localized lipid peroxidation"

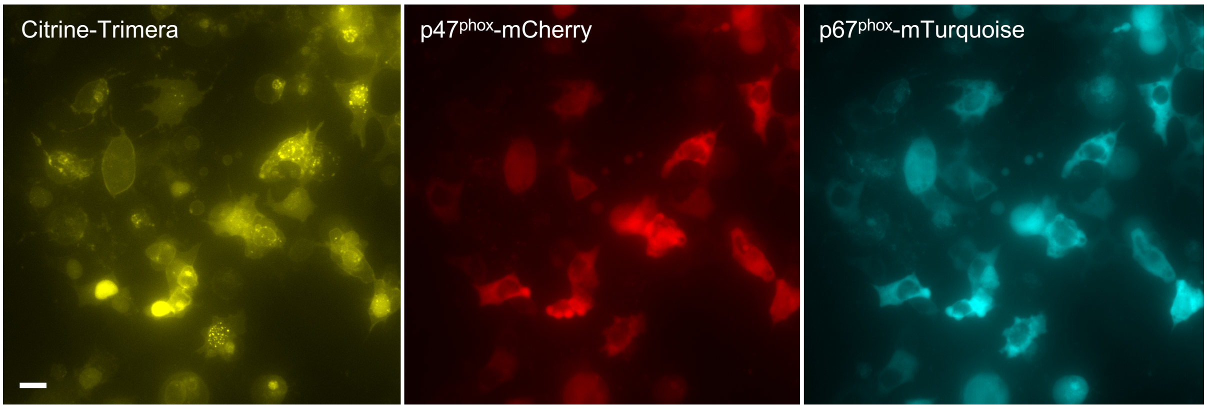


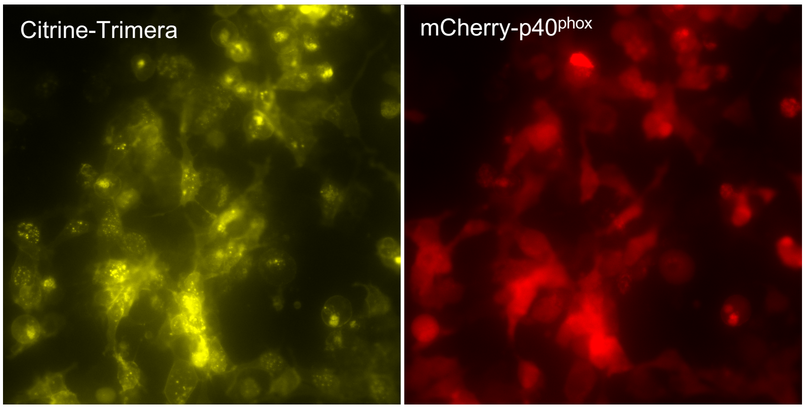


**Transfected cells (%)**

| Condition | CFP | YFP | RFP |
|---|---|---|---|
| Citr-Trimera + p47-mCherry + p67-mTurq | 70 | 80 | 65 |
| Citr-Trimera + mCherry-p40 | | 87 | 82 |
| p47-mCherry + p67-mTurq | 72 | | 75 |

***Figure S1. Transfection efficiency in COSNOX cells transfected with Citrine-Trimera and/or FP-labeled cytosolic subunits.*** *COSNOX cells transfected with Citrine-Trimera, p47$^{phox}$mCherry and p67$^{phox}$mTurquoise (top panel) or with Citrine-Trimera and mCherryp40$^{phox}$ (bottom panel). Table on the right shows transfection efficiency for cells in corresponding images, and for COSNOX cells transfected only with separated subunits (p47$^{phox}$mCherry and p67$^{phox}$mTurquoise (images not shown). Images acquired by wide-field microscopy. Scale bar 10 µm.*

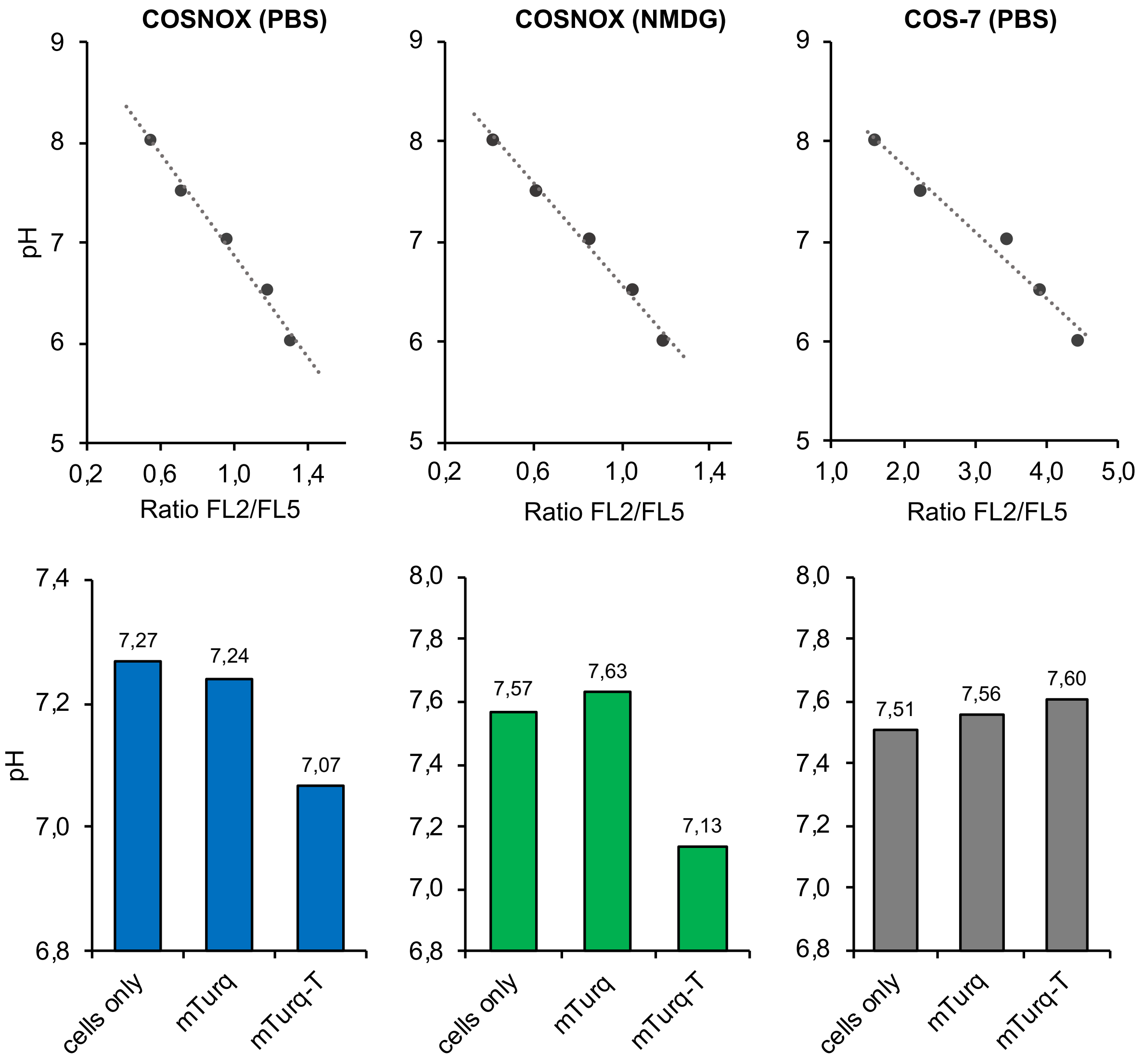


***Figure S2. Top:*** *Representative calibration curves of SNARF-1 probe. Ratio FL2/FL5 means ratio of the fluorescence intensity of the protonated/deprotonated form of the SNARF-1 probe. FL2 and FL5 are custom names according to the channels of the flow cytometer in our laboratory. COSNOX cells were investigated either in classic PBS buffer or in $Na^+$ free buffer (NMDG). COS-7 cells were used as a negative control, because they do not contain any of the membrane subunits of the NADPH oxidase, so they are unable to produce ROS in presence of the Trimera protein.* ***Bottom:*** *Intracellular pH in COSNOX and COS-7 24h after transfection by mTurquoise-Trimera or mTurquoise calculated using the calibration curves showed above.*

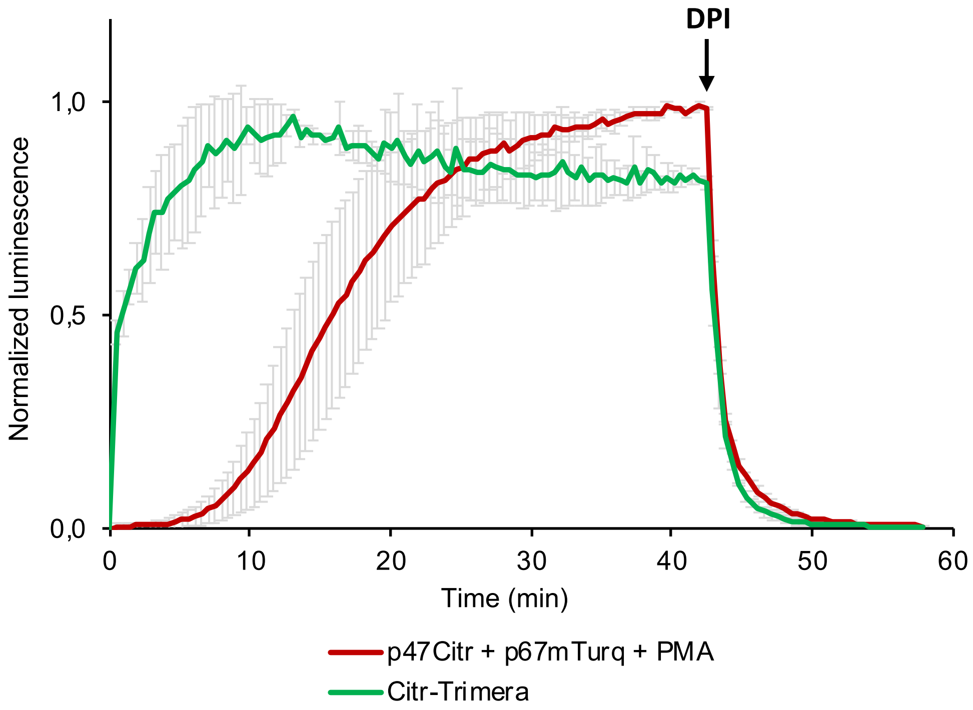


***Figure S3. Trimera triggers continuous NADPH oxidase activity in cells.*** *Normalized ROS production detected by luminometry in COSNOX cells transfected with Citrine-Trimera or with p47$^{phox}$mCherry and p67$^{phox}$mTurquoise. Cells transfected with individual cytosolic subunits were activated by 1 µM PMA. The DPI (25 µM) was added 42 min after the start of the measurement. Error bars show SD for n = 4 experiments.*

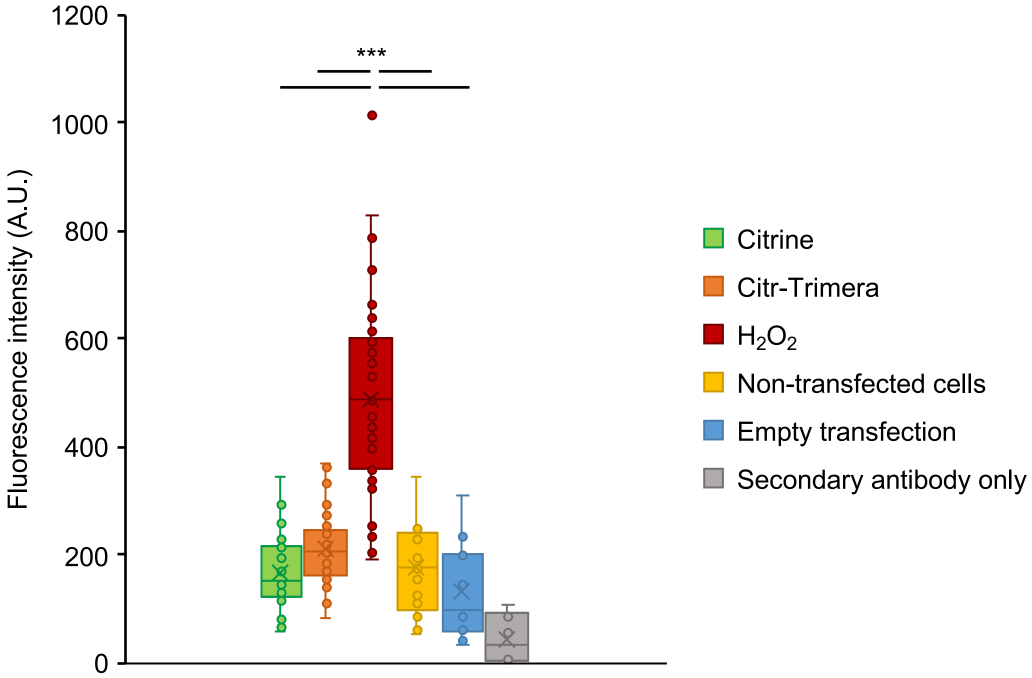


***Figure S4. Levels of 4-HNE in COSNOX cells 24h after transfection with Citrine-Trimera.*** *Fluorescence intensity of Alexa Fluor 647 (coupled to the secondary antibody) corresponds to the amount of the 4-HNE in analyzed COSNOX cells. As a positive control, COSNOX cells were treated with 200 µm $H_2O_2$ overnight. Statistical analysis performed by one-way ANOVA followed by a Tukey's Multiple Comparison Test (*** means $p < 0.001$). Data from 2 independent experiments.*

### Calibration of the wide-field microscope for FRET-FLIM experiments

In wide-field microscopy, the observation volume is determined as the product of the surface of the ROI and the cell thickness (height). Due to the biological diversity, each cell has a different thickness, which makes the determination of a generic cell volume impossible. So, the intracellular concentration of a single FP cannot be calculated directly from the fluorescent intensities. We overcome this issue by a ratiometric approach [1]. To determine the ratio of the amount of the acceptor to the donor in cells, a calibration of the fluorescence intensities for the donor and the acceptor was performed using droplets of purified fluorescent proteins. In living cells, the relative expression level of fluorescent proteins, which can be described as a ratio of the amounts of the acceptor to the donor ($n_A/n_D$), is proportional to the ratio of their fluorescence intensities ($n_A/n_D \propto I_A/I_D$). Supposing that the observation volume is identical for the donor and the acceptor in the cell, as it is the case for cytosolic proteins, we can rewrite this relation using donor and acceptor concentrations: $[A]/[D] \propto I_A/I_D$. The coefficient of proportionality, called calibration factor $f$, between $[A]/[D]$ and $I_A/I_D$ was evaluated using 50 µL droplets of solutions of purified fluorescent proteins. In those droplets, the observation volume is entirely contained in the droplet and can be simplified.

The purified FPs, mTurquoise and Citrine, with known concentration were diluted in series. A droplet (50 µL) of a diluted solution was placed on a glass cover slip in a Attofluor chamber, covered by another cover slip to avoid evaporation, which would cause an artificial increase of the protein concentration. Images were acquired and the mean fluorescence intensity of the whole field of view was normalized and plotted against the FP concentration. The data points of each protein were fitted with a linear fit function with an intercept at zero (Figure S5).

The values of the slopes of the trendlines for mTurquoise and Citrine were then used to determine the calibration factor *f*:

$$\frac{[A]}{[D]} = \frac{[YFP]}{[CFP]} = \frac{I_{YFP}}{I_{CFP}} f = \frac{I_{YFP}}{I_{CFP}} \frac{slope\ (CFP)}{slope\ (YFP)} = \frac{I_{YFP}}{I_{CFP}} 1.35 \qquad \textit{(Equation S1)}$$

$$f = 1.35$$

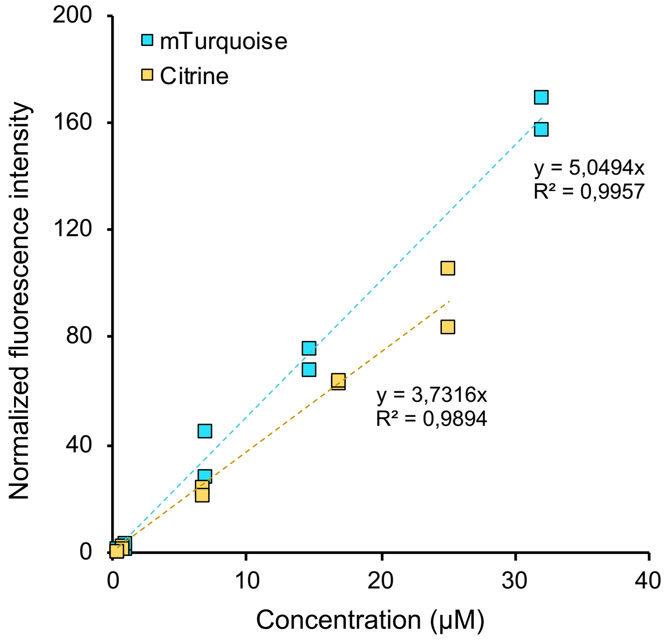


***Figure S5.*** *Calibration curves of mTurquoise and Citrine.*

A

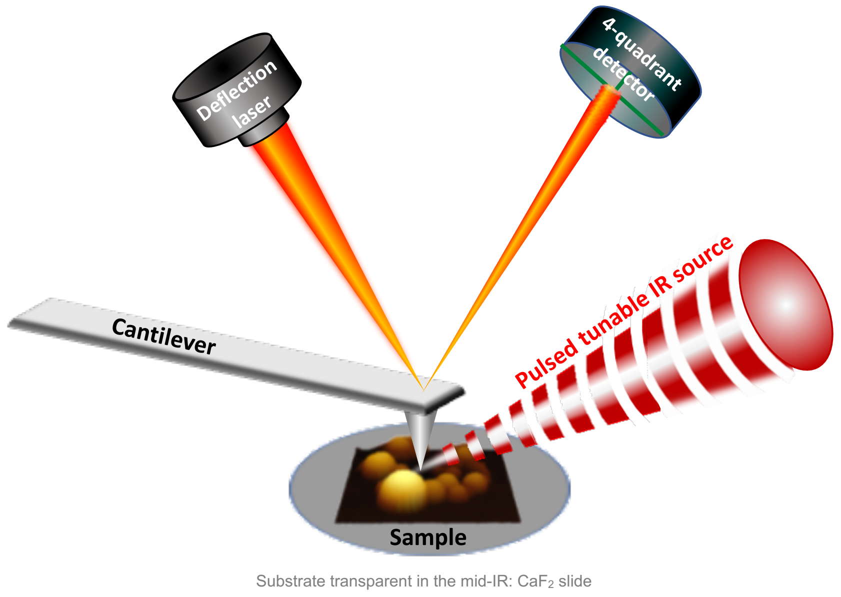


B

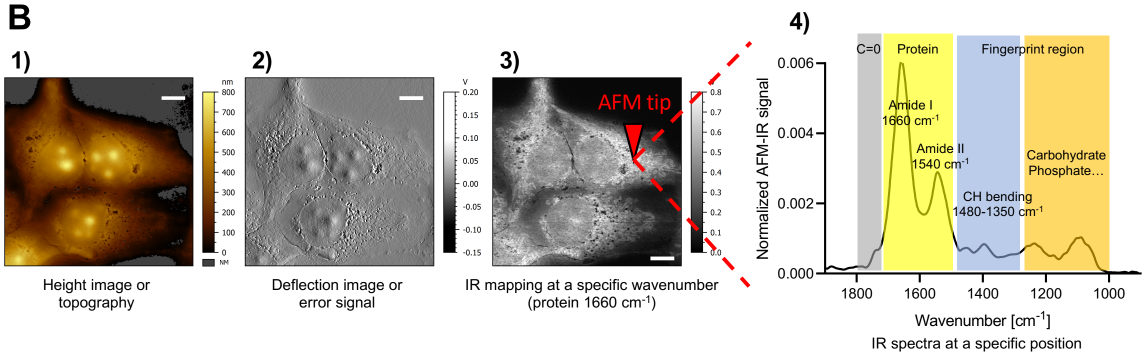


***Figure S6. AFM-IR technology. A:*** *A pulsed tunable laser source is focused on a sample near the AFM cantilever tip. When the laser is tuned to an absorbance band of the sample, the absorbed light causes a short-lived photothermal expansion, which leads to resonant oscillation of the AFM tip. The oscillations are detected by the deflection laser position on the 4-quadrant detector. The amplitude of the tip oscillation is directly proportional to the sample absorption coefficient. In our study, we used resonance-enhanced AFM-IR that improves the sensitivity compared to the conventional AFM-IR by pulsing the laser at a frequency matching the resonance frequency of the AFM cantilever.* ***B:*** *Using AFM-IR we can obtain height image (B1), deflection image (B2) or IR map at a specific wavenumber (B3) of a large area of the sample. Local absorption spectrum (B4) provides information about chemical composition at the concrete acquisition site.*
